# Automated and modular protein binder design with BinderFlow

**DOI:** 10.1101/2025.09.10.675490

**Authors:** Carlos Chacón-Sánchez, Nayim González-Rodríguez, Oscar Llorca, Rafael Fernández-Leiro

**Affiliations:** Structural Biology Programme, Spanish National Cancer Research Centre (CNIO), Madrid, Spain

## Abstract

Deep learning has revolutionised *de novo* protein design, with new models achieving unprecedented success in creating novel proteins with specific functions, including artificial protein binders. However, current methods remain computationally demanding and challenging to operate without specialised infrastructure and expertise. To overcome these limitations, we developed `BinderFlow`, a structured and parallelised pipeline for protein binder design. Its batch-based nature enables live monitoring of design campaigns, seamless coexistence with other GPU-intensive processes, and reduces human intervention. Furthermore, `BinderFlow`’s modular structure enables straightforward modifications to the design pipeline to incorporate new models and tools or to implement alternative design strategies. Complementing this, we developed `BFmonitor`, a web-based dashboard that simplifies campaign monitoring, design evaluation, and hit selection. Together, these tools lower the entry barrier for non-specialised users and streamline expert workflows, making generative protein design more accessible, scalable and practical for both exploratory and production-level research.

---

Proteins are complex biomolecules that perform many different functions, from catalysing chemical reactions to modulating regulatory networks through protein-protein interactions (PPIs). This functional versatility is mediated through the wide range of structures they adopt^1^. However, the structural space explored by natural proteins is limited, and thus the functions these proteins fulfil do not represent the complete range of potential tasks^2,3^. The possibility of creating proteins from scratch^4^ with tailored functions^5^ has made *de novo* protein design a major goal in molecular biology.

Sampling the protein-sequence space using biophysical methods quickly becomes computationally intractable due to its vast dimensions. The recent development of generative models provides an alternative to physics-based methods. These deep-learning models, trained on large datasets of proteins, can generate structures and sequences that fulfil a pre-specified function from scratch using a moderate amount of resources and time^6–9^.

One of the many applications of artificial proteins is to bind to a specific surface of interest on a target protein, modulating its functions^10–13^. Traditionally, the production of such PPI regulators has required animal immunisation for antibody production or massive screening of chemical libraries. These methods require considerable experimental effort and present certain limitations, mainly: some interfaces lack defined pockets for small molecule targeting^14,15^, antibody development involves high production and conservation costs^16^, and some targets are not suitable for chemical screening^13,15^. Artificial proteins designed specifically to bind to a protein interface —binders from now on— overcome these problems, as they can be produced in *Escherichia coli*, feature extreme thermal stability, and exhibit high affinity and specificity towards their target^11,17,18^. Thus, the possibility to engineer binders in a targeted manner has both immediate therapeutic^12,19^ and biotechnological implications^20,21^.

A typical protein binder design project begins by selecting the region of interest in the surface of the target, constraining the design space. The process typically involves finding a backbone whose shape is complementary to the target surface and assigning a sequence of amino acids that folds into that backbone to establish intermolecular interactions with the target. The pipeline described by Watson *et al*.^22^ has become the standard for *de novo* binder generation^18^. This pipeline uses Rosetta Fold Diffusion (RFD), a diffusion model that “denoises” previously unseen backbones from randomly distributed atoms^22^, ProteinMPNN (pMPNN) to assign sequences to those backbones^23^, and a modified version of AlphaFold2 (AF2IG) to assess the quality of the designs *in silico*^24–26^, sequentially. AF2IG confidence metrics are then used to filter *in silico*-successful binders —hits from now on— which are then validated experimentally (Fig. 1A).

**Fig. 1.**
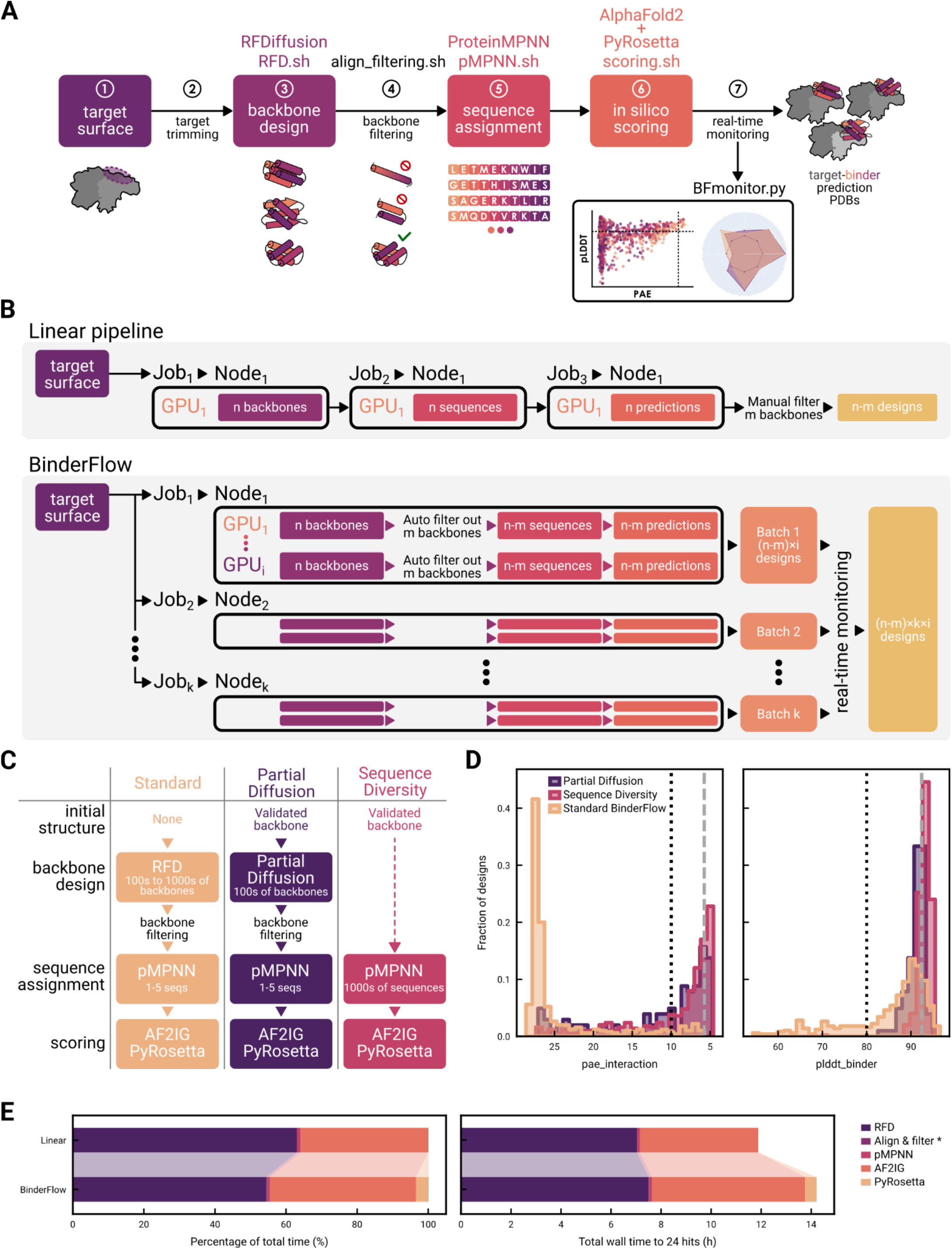
BinderFlow pipeline: architecture and alternative strategies. **A)** Schematic of the BinderFlow pipeline: 1. The user defines a hotspot in a surface of interest. 2. The target structure gets trimmed to increase computational e=iciency. 3. RFD.sh produces protein backbones of a specified length complementary in shape to the target. 4. align_filtering.sh filters out suboptimal backbones that might be problematic for expression, such as long helices or isolated hairpins. 5. pMPNN.sh assigns a sequence of amino acids to each backbone. 6. scoring.sh predicts the binder-target complex using AF2IG, collects the relevant AF2 scores and uses PyRosetta for measuring a set of relevant parameters for interactions. 7. The process is monitored in real time using the BFmonitor.py web-based tool. **B)** Architecture of BinderFlow and comparison with a linear workflow. In a typical workflow each step of the pipeline requires manual handling of input and output files, and manual inspection to remove obvious suboptimal candidate backbones. Moreover, each step is run as an independent job, hampering parallelisation. Meanwhile, BinderFlow distributes batches of end-to-end predictions as independent jobs, facilitating parallelisation of instances across GPUs and HPC cluster nodes. It includes automatic filtering of suboptimal backbones and real-time monitoring of the process, allowing for stopping the campaign once a suitable number of hits is achieved. **C)** Comparison between the standard BinderFlow pipeline and proposed alternative strategies for binder refinement. The dashed arrow indicates skipping the backbone generation step. **D)** Comparison of PAE interaction and binder pLDDT score distributions resulting from the standard BinderFlow pipeline, Partial Di=usion and Sequence Diversity. The grey, dashed lines indicate the pae_interaction and plddt_binder scores of the design used to initialize Partial Di=usion and Sequence Diversity runs, obtained from the standard BinderFlow pipeline. The black, dotted lines indicate typical thresholds to consider a binder a hit, PAE_interaction < 10 and pLDDT_binder > 80. **E)** Comparison of time spent per step using a linear binder design pipeline and the BinderFlow architecture. Left: normalized times and proportion of the run length allocated to each step. Right: comparison of the total wall times required to obtain 24 hits. *Align C filter times are too small to be noticed in the bar plot (Figure S1).

As structural prediction scores are not infallible indicators of actual binding, many designs still need to be screened and validated experimentally to find a successful one. Typically, tens or hundreds of protein designs are tested on plate-based assays for convenience and throughput, previously selected from computational campaigns that range from thousands to tens of thousands of candidates^10,12,17,22,27^. Such campaigns require computational resources that are often prohibitive for small or non-specialised laboratories. In addition, the default way to execute this pipeline^11,22,28^ is to run each step sequentially as single, large jobs in a linear fashion, i.e., generating thousands of backbones, then calculating the corresponding thousands of sequences, and finally evaluating thousands of potential binders using AF2IG. Notably, the *in silico*-success rate for a given campaign, meaning the ratio between hits and total designs, varies widely^11,17,22^ and cannot be estimated beforehand. This often results in inefficient campaigns, where the number of needed designs is either overestimated, wasting limited computational resources, or underestimated, forcing the launch of a new set of jobs to increase the total size of the campaign.

Here, we report `BinderFlow,` a pipeline designed to make protein binder design more efficient and accessible, democratising *de novo* protein binder design for the wider scientific community. This approach divides a typical protein design project into arbitrarily sized batches, each performing backbone design, sequence assignment and candidate scoring in an automated workflow. By breaking the campaign into shorter tasks, `Binder-Flow` enables more granular execution and live monitoring of progress. This structure also enables researchers to opportunistically use GPU resources —usually employed by unrelated workflows— for executing short-lived protein design jobs when idle. Compared to a linear approach, `BinderFlow` facilitates binder design through parallelisation and live monitoring, and by preventing superfluous calculation of more hits than can be experimentally screened. Overall, `BinderFlow` makes protein design more accessible, enabling its coexistence with other work in the same computing infrastructure.

We provide fully integrated code for deploying `BinderFlow` on SLURM-based^29^ HPC clusters or smaller setups, such as individual workstations, alongside `BFmonitor`, a web-based application from which the user can interact with the pipeline, monitor the campaign in real time and prepare candidates for down-stream applications.

## The BinderFlow pipeline

In `BinderFlow`, we have integrated backbone design, filtering of suboptimal backbones, sequence inference and score calculations in a continuous workflow, automating the input and output handling between each step (Fig. 1A). As a result, binder design can be executed end-to-end in batches of a small number of designs per job.

A design campaign is split into multiple, parallel `BinderFlow` instances, each addressing a single batch of designs and independently executed on available GPUs (Fig. 1B). Each `Binder-Flow` instance executes the following scripts sequentially per GPU (Fig. 1A):

1. `RFD.sh:` runs RFD to produce *n* binder backbones.
2. `align_filtering.sh:` replaces the cropped target chain with a complete version, providing more context for the sequence assignment and scoring algorithms. To avoid subsequent waste of computational resources, it also filters out designs with steric clashes and backbones composed of a single, long helix or a single hairpin, which are difficult to produce experimentally.
3. `pMPNN.sh:` runs pMPNN to assign sequences to the binder backbone in the context of the target.
4. `scoring.sh:` predicts the binder-target complex structure using a modified version of AF2IG^24–26^ to obtain the per-residue predicted Local Distance Difference Test (pLDDT) and Predicted Aligned Error (PAE) scores. Then, it calculates physics-based metrics using PyRosetta^30^ to further filter and characterise the designs (e.g., shape complementarity^31^ or number of unsatisfied hydrogen bonds^32^). Before finishing, every`BinderFlow` instance appends the scores associated with each design to a .*csv* file for live monitoring of the campaign (Fig. 1A) and evaluates the number of designs that meet the *in silico* conditions for a binder to be considered a hit (e.g., PAE_interaction < 10, pLDDT_binder > 80). This workflow is then repeated until the desired number of hits is obtained. This architecture is fully modular, so each step can be adapted to a different software that performs similar functions, and further steps can be added or substituted as new tools become available.

The end-to-end architecture of `BinderFlow` instances allows independent execution and, thus, their parallelisation across different GPUs (Fig. 1B). Additionally, `BinderFlow` jobs can be submitted with low priority to the queuing system, ensuring they are executed only when GPU nodes would otherwise remain idle. This facilitates the simultaneous execution of multiple campaigns and the parallelisation of multiple GPUs within a single campaign, reducing HPC queue clogging and improving the efficiency of shared computational resources. Overall, this workflow makes binder design more accessible to researchers with access to GPU infrastructure, but not primarily focused on protein design, such as structural biology laboratories.

## BinderFlow modularity enables binder refining strategies

Once an artificial or natural binder has been validated experimentally, it might still not meet desired biochemical criteria (e.g., affinity, specificity, solubility). Using protein design tools, it is possible to enhance their properties by further exploring the structural and sequence spaces. To improve previously identified binders, we have implemented two refining strategies into our pipeline: Partial Diffusion^22^ and Sequence Diversity.

Partial Diffusion employs a method previously described by Vazquez-Torres *et al*.^21^, in which, instead of random coordinates, the atomic positions of a validated backbone are used as the starting point. These positions are disturbed by adding white noise —typically a fraction of the noise level used for RFD— and then “denoised” to explore the nearby structural space, where new energy minima might be found. This step replaces regular RFD backbone inference, and the rest of the process follows the same steps as the pipeline described above (Fig. 1C).

The Sequence Diversity strategy uses pMPNN to sample the sequence space. pMPNN designs new sequences that likely fold into the previously specified backbone, while integrating the context of the target interface. Providing a validated backbone as input, we massively run pMPNN to obtain many sequences that are predicted to fold into the same structure (Fig. 1C). This strategy fully skips the backbone generation step, making it less computationally expensive than Partial Diffusion.

Compared to *de novo* backbone design, the distribution of *in silico* metrics after Partial Diffusion is skewed towards what is considered a hit (Fig. 1D, Supplementary Fig. 1A). The improvement in metrics is likely due to a better fit of the backbones to the target surface (Supplementary Fig. 1B, 2A). The distribution of designs from Sequence Diversity changed similarly (Fig. 1D, Supplementary Fig. 1A). The increase in success rate correlates with the shape complementarity of the binder sequence to the target (Supplementary Fig. 1C), but not with sequence similarity to the original sequence (Supplementary Fig. 1D,E). This demonstrates that Sequence Diversity gently explores the structural space around the input structure, as different sequences induce slight structural adjustments to accommodate the different amino acids while maintaining the overall fold (Supplementary Fig. 2). Further structural exploration within Sequence Diversity could be achieved by performing pMPNN with FastRelax, a protocol that slightly modifies the binder backbone by energy minimisation^24,33^. However, we have not observed an increase in *in silico* hits obtained using FastRelax on RFD-calculated backbones (Supplementary Fig. 3), as others have noted^22^.

**Fig. 2.**
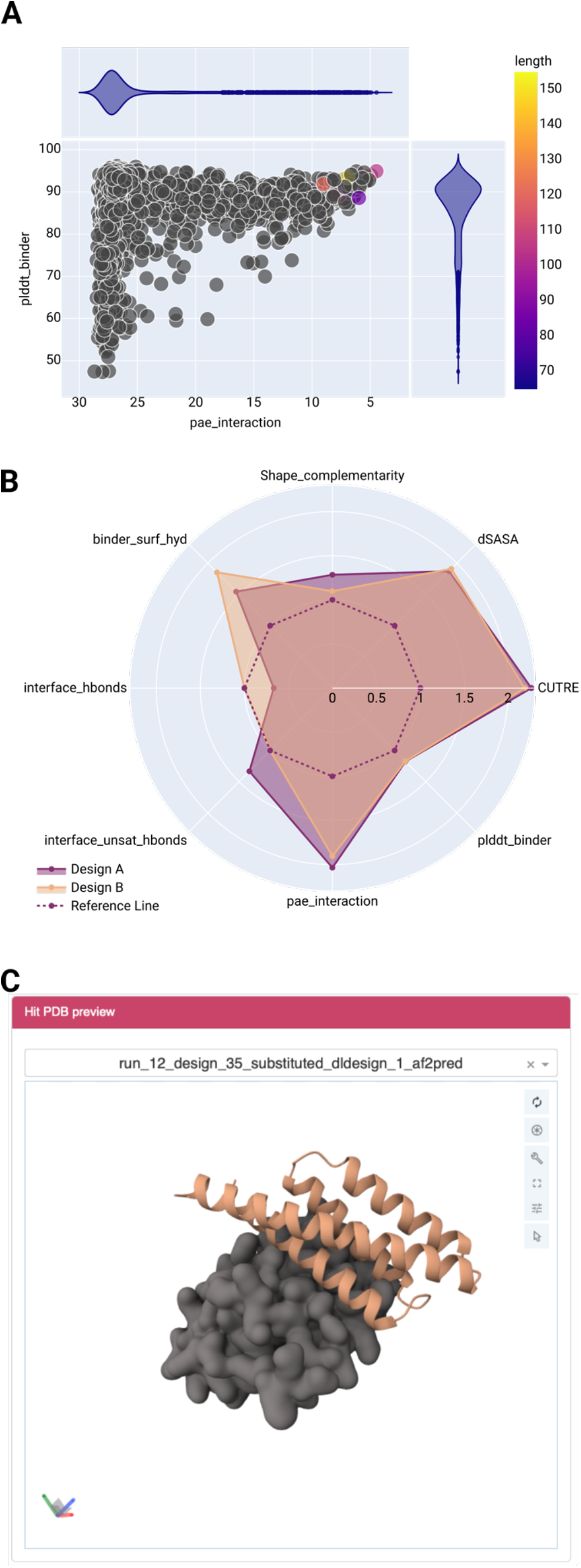
The BFmonitor web-based dashboard. BFmonitor provides three tools to follow a design campaign in real time (see Fig. S4 for a full view of the dashboard). **A)** A scatter plot that updates in real time, providing information about any two of the variables calculated by *scoring*.*sh*. Each data point corresponds to an individual binder, and only those that fulfil all *in silico* criteria get colored. **B)** A radar plot enables pairwise comparison of all scores calculated for any two binders, providing visual information on how each metric compares between the two candidates. **C)** An interactive 3D viewer allows to inspect the backbones of *in silico* hits (yellow cartoon) interactively in the context of the target protein (grey surface) in real time.

It is important to note that the performance of each design strategy will vary across proteins, as different backbones require different degrees of exploration of the sequence and structural landscape to optimise binding to their targeted surface. BinderFlow enables the efficient exploration and combination of multiple design strategies, facilitating the design process.

## BinderFlow benchmarking

To characterise how the batch-based architecture of `Binder-Flow` affects the performance of the pipeline, we launched a binder design campaign against Programmed Death-Ligand (PDL1), a protein often employed as a benchmark for binder generation^17,22,34^. The designs ranged from 65 to 155 residues (Supplementary Fig. 2A), using both `BinderFlow` and the linear pipeline (see Methods for details). The efficiencies of RFD and pMPNN remained largely unchanged. The linear pipeline averages 63.5 s and 1.0 s per design, respectively, compared to 68.0 s and 1.1 s per design in `BinderFlow` (Fig. 1E, Supplementary Fig. 4A). Initially, the total processing time per binder increased due to the AF2IG scoring step, which required 273% more time when utilising the `BinderFlow` structure (134.3 s using `BinderFlow` versus 36.1 s using the linear pipeline). Splitting large jobs into smaller batches required restarting calculations for each batch, which came at the cost of losing intermediate results stored in memory. Upon inspecting the AF2IG code^24^, we observed that each prediction required an intermediate calculation that is computed once for a given design length and then recycled for all designs with the same number of amino acids, reducing scoring times by 10-fold (Supplementary Fig. 4B). By splitting the design campaign into batches, this calculation took place hundreds of times instead of once per design length, significantly slowing down the process. We alleviated this inefficiency by writing the matrices resulting from those calculations to disk and reusing them across batches. Thus, the modified version of AF2IG requires only 51.3 s per binder when executed as part of `BinderFlow`, reducing the total time difference between the linear and `BinderFlow` pipelines to 24% (Fig. 1E, Supplementary Fig. 4A,C). Importantly, the time lost per binder due to slower scoring was partially offset by automatic backbone filtering after RFD (Fig. 1A), which avoids wasting time on sequence inference and scoring of suboptimal backbones. It took `BinderFlow` 14.2 h of total wall time to obtain 24 hits, compared with 12 h for the linear pipeline. Thus, the difference in efficiency is reduced from 24% to 18% (Fig. 1E, Supplementary Fig. 4A,C). Note that this comparison assumes the best-case scenario for the linear pipeline, where the run’s efficiency is known, thereby avoiding the waste of computation on more hits than required and the time spent re-running the pipeline due to underestimation. In real-case scenarios, `BinderFlow`‘s live monitoring of the hit rate prevents superfluous calculation of more hits that can be experimentally screened, reducing even further the efficiency gap.

## Real-time monitoring

Another advantage of `BinderFlow` and its end-to-end binder design architecture is that it enables real-time monitoring of design campaigns as scores are calculated in small batches. We have developed `BFmonitor`, a web-based dashboard that includes tools to monitor campaigns, evaluate designs, and select hits for DNA synthesis (Fig. 2, Supplementary Fig. 5).

The first tool is `Live Watcher`, an interactive graphical summary of the design campaign. In it, the user can set thresholds for all parameters calculated by AF2IG and PyRosetta to define when binders are considered hits and filter them. It contains a scatter plot, in which any pair of these parameters can be represented, providing an overview of the project and identifying correlations between them (Fig. 2A, Supplementary Fig. 5A). It also includes a radar plot for pairwise comparisons of designs, showing all calculated parameters normalised by the value set for the mentioned thresholds (Fig. 2B, Supplementary Fig. 5A). The second tab, `Pipeline tracking`, details the progress of each instance. The last tab, `Extraction` (Fig. 2C, Supplementary Fig. 5B), allows the user to preview the structure of the binder-target complex for any design that passes the thresholds. In this window, the user can extract the hits structures as. `pdb` files and their sequences in `.fasta` format. In addition, we include a reverse translation tool based on CodonTransformer^35^, which returns DNA coding sequences for the selected binders. These can be flanked by custom 5’ and 3’ sequences to facilitate cloning and downstream applications, and can be directly used for DNA synthesis orders.

## Discussion

The introduction of generative models has significantly streamlined the process of binder design. Tasks that previously required substantial expertise in protein engineering can now be executed efficiently using these computational tools. As a result, the main bottleneck has shifted from technical knowledge in protein design to biological insights for selecting relevant targets and robust validation strategies. Yet, while many laboratories have the necessary biological expertise, they often lack the computational infrastructure and expertise required to deploy large-scale binder design projects. Moreover, repurposing existing computational resources to support protein design is not always feasible.

Here, we introduce `BinderFlow`, a pipeline designed to democratise and facilitate *de novo* protein binder design for both non-experts and experts. `BinderFlow` divides otherwise large protein design campaigns into batches, enabling their coexistence with other resource-intensive activities in the same computing infrastructure. It features a parallelizable architecture and automates most tasks that usually require human intervention, such as handling input and output or rejecting suboptimal candidate backbones. Its batch-based design enables live monitoring of the campaigns, allowing for the estimation of campaign efficiency in real-time. Moreover, computational resources are allocated efficiently by avoiding superfluous calculations after the desired number of hits is obtained.

We provide `BinderFlow` as a ready-to-use implementation for SLURM-based^29^ systems, ranging from HPCs to individual workstations. Although the GPU time per binder is increased using this pipeline compared to a linear workflow (Fig. 1E), this difference can be alleviated by increasing the batch size and reducing the length range (Supplementary Fig. 4B), albeit at the cost of a lower frequency of monitoring updates.

To keep pace with the rapid advances of the protein design field, we designed `BinderFlow` as a modular pipeline, streamlining the incorporation of new software to give users the option to use their preferred tool for each step. This modularity also enables flexible adaptation of the design workflow itself. We illustrate this by providing two ready-to-use strategies for binder refinement that explore both the structure and sequence space: Partial Diffusion and Sequence Diversity. Our *in silico* results indicate that these are promising strategies to improve binder affinity and specificity. Nonetheless, the relationship between computational confidence metrics and experimental binder success rates remains poorly understood. Advancing the experimental efficiency of protein design will require new, more predictive metrics that integrate quantitative affinity or phenotypic data collected by coherent experimental approaches, prediction models, and biophysical scoring functions.

different GPUs (Fig. 1B). Additionally, `BinderFlow` jobs can be submitted with low priority to the queuing system, ensuring they are executed only when GPU nodes would otherwise remain idle. This facilitates the simultaneous execution of multiple campaigns and the parallelisation of multiple GPUs within a single campaign, reducing HPC queue clogging and improving the efficiency of shared computational resources. Overall, this work-flow makes binder design more accessible to researchers with access to GPU infrastructure, but not primarily focused on protein design, such as structural biology laboratories.

### BinderFlow modularity enables binder refining strategies

Once an artificial or natural binder has been validated experimentally, it might still not meet desired biochemical criteria (e.g., affinity, specificity, solubility). Using protein design tools, it is possible to enhance their properties by further exploring the structural and sequence spaces. To improve previously identified binders, we have implemented two refining strategies into our pipeline: Partial Diffusion^22^ and Sequence Diversity.

Partial Diffusion employs a method previously described by Vazquez-Torres *et al*.^21^, in which, instead of random coordinates, the atomic positions of a validated backbone are used as the starting point. These positions are disturbed by adding white noise —typically a fraction of the noise level used for RFD— and then “denoised” to explore the nearby structural space, where new energy minima might be found. This step replaces regular RFD backbone inference, and the rest of the process follows the same steps as the pipeline described above (Fig. 1C).

The Sequence Diversity strategy uses pMPNN to sample the sequence space. pMPNN designs new sequences that likely fold into the previously specified backbone, while integrating the context of the target interface. Providing a validated backbone as input, we massively run pMPNN to obtain many sequences that are predicted to fold into the same structure (Fig. 1C). This strategy fully skips the backbone generation step, making it less computationally expensive than Partial Diffusion.

Compared to *de novo* backbone design, the distribution of *in silico* metrics after Partial Diffusion is skewed towards what is considered a hit (Fig. 1D, Supplementary Fig. 1A). The improvement in metrics is likely due to a better fit of the backbones to the target surface (Supplementary Fig. 1B, 2A). The distribution of designs from Sequence Diversity changed similarly (Fig. 1D, Supplementary Fig. 1A). The increase in success rate correlates with the shape complementarity of the binder sequence to the target (Supplementary Fig. 1C), but not with sequence similarity to the original sequence (Supplementary Fig. 1D,E). This demonstrates that Sequence Diversity gently explores the structural space around the input structure, as different sequences induce slight structural adjustments to accommodate the different amino acids while maintaining the overall fold (Supplementary Fig. 2). Further structural exploration within Sequence Diversity could be achieved by performing pMPNN with FastRelax, a protocol that slightly modifies the binder backbone by energy minimisation^24,33^. However, we have not observed an increase in *in silico*

## Methods

### Benchmarking

For all benchmarking campaigns, consumer graphics cards (RTX2080Ti or RTX4090Ti) were employed as indicated, as they represent standard and affordable cards commonly used in laboratories employing GPU-based routines with relatively low memory requirements, such as structural biology groups.

For the `BinderFlow` benchmarking campaigns, a trimmed version of PDL1 (residues 18-132, AFDB AF-Q9NZQ7-F1-v4) was used as target to decrease computation times. The selection of this protein is due to its use as a benchmark in previous publications with different binder generation models^17,22,34^. The structure was renumbered to start in chain B and residue 1018 to prevent indexing issues with AF2IG. Residues 1054 and 1068 (I54 and V68 in untrimmed PDL1, respectively) were selected as hotspots for binder generation. Binders of lengths comprising 65-155 residues were designed using RFD, with default parameters (complex_base checkpoint, 50 noise steps). 10-backbone batches were designed per available GPU.

Binder backbones were selected attending to: (1) the presence of steric clashes (defined as having at least one atom at less than 0.5 Å from the target structure), and (2) their tertiary structure, filtering out designs predicted to fold as harpins or long, single helices using DSSP^36^. One sequence per backbone was generated using pMPNN without FastRelax optimization^24^, and binder-target complexes were used as input for AF2IG scoring and PyRosetta analysis to extract different biophysical metrics (difference in Solvent Accessible Surface Area, interface hydrogen bonds, unsatisfied interface hydrogen bonds^32^ and binder surface hydrophobicity. The campaigns were run until obtaining at least 48 *in silico* hits.

For the campaigns following the linear pipeline, the same input and hotspots were used. For the backbone generation step, 520 and 680 designs were generated using RFD with the RTX 2080 Ti and RTX 4090 Ti GPUs, respectively, based on our previous experience with PDL1 binder design hit rate. Next, we generated one sequence per design using pMPNN without FastRelax and, lastly, they were scored using AF2IG.

A binder is considered a hit following the description by Bennet et al.^24^ and followed by others^17,22,37,38^ (pLDDT_binder > 80 and PAE_interaction < 10). Other scores, included in PyRosetta^30^, or derived from AF2 scores such as ipSAE^39^, are also implemented to further filter *in silico* hits.

### Sequence diversity and partial diffusion runs

Partial Diffusion and Sequence Diversity campaigns following `BinderFlow` architecture were performed only in Nvidia RTX2080Ti GPU cards, as we did not observe a significant change in efficiency with respect to the RTX4090Ti GPU cards. Five independent candidate binders were selected from the benchmarking campaigns as input for both Sequence Diversity and Partial Diffusion, based on their metrics and the presence of viable, diverse folds (Fig. S3). For Partial Diffusion campaigns, default noise settings were employed (Complex_base as checkpoint, 20 noising steps, noise scale of 1 for both translations and rotations). 10-backbone batches were designed per available GPU. Backbones were filtered, assigned a single sequence using pMPNN without Fast Relax and scored by AF2IG and PyRosetta as indicated above. The campaign continued until at least 48 hits were obtained.

For Sequence Diversity campaigns, between 5000 and 5400 sequences were designed in batches of 200 sequences per GPU using pMPNN without the Fast Relax protocol. The designs were then scored using AF2IG and PyRosetta. The same thresholds for hits as in previous campaigns were used.

For Sequence Diversity with Fast Relax campaigns, 200 sequences were designed per input structure (same as in Partial Diffusion and Sequence Diversity campaigns), in batches of 1 sequence with 1 Fast Relax cycle. Designs were scored as indicated above.

### Implementation

`BFmonitor` is written in Python and uses Dash from Plotly^40^ for its deployment as a web-based application. All the information is stored in `.csv` files and processed using the pandas package^41^. Both `BinderFlow` and `BFmonitor` are currently compatible with RosettaFold Diffusion and pMPNN, but the modular organization of the scripts make it easily adaptable to other open-source tools. Installation, usage instructions and tutorials for these tools are available at https://github.com/cryoEMCNIO/BinderFlow.

## Data availability

PDL1 was used as a case study. Its sequence and structure prediction are available at AFDB AF-Q9NZQ7-F1-v4.

## Code availability

The code for `BinderFlow` and `BFmonitor` is publicly available at https://github.com/cryoEM-CNIO/BinderFlow.

## Acknowledgements

We thank members of CNIO Structural Biology Programme for their critical feedback and for beta testing the pipeline. We are also grateful to our colleagues for their constructive reading and comments on the manuscript.

## Author contributions

C.C.-S.: Methodology, Software, Investigation, Writing – original draft, Writing – review & editing. N.G.-R.: Conceptualization, Methodology, Software, Investigation, Writing – original draft, Writing – review & editing. O.L.: Conceptualization, Writing – review C editing. R.F.-L.: Conceptualization, Methodology, Software, Project administration, Writing – original draft, Writing – review C editing, Supervision, Corresponding author.

