## supplementary figures for "Automated and modular protein binder design with BinderFlow"

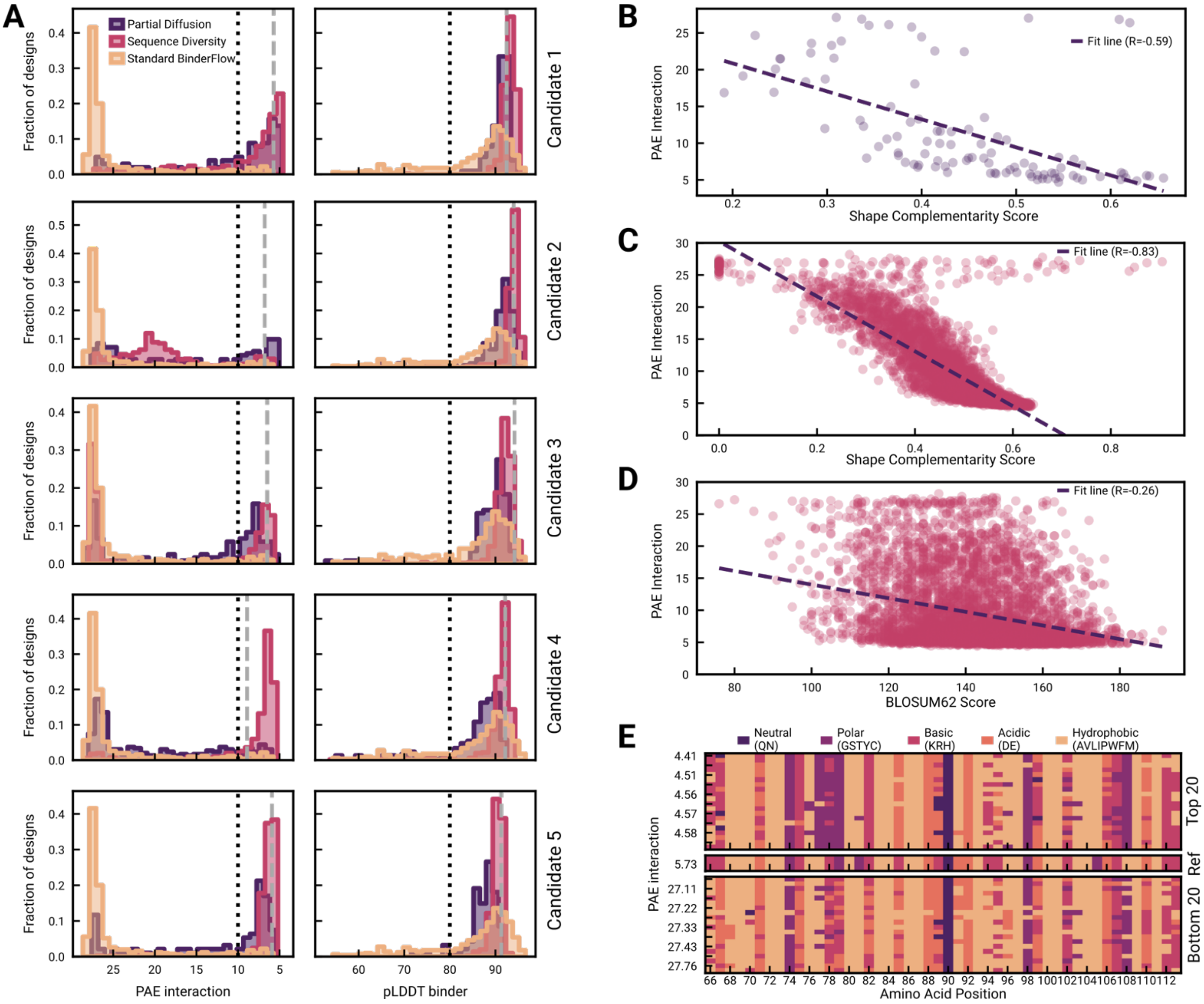


**Supplementary Fig. 1 – Efficiency comparison between the standard, Partial Diffusion and Sequence Diversity pipeline**

A) Comparison of PAE interaction and binder pLDDT distributions resulting from campaigns run using the standard BinderFlow pipeline, Partial Diffusion and Sequence Diversity. Each row represents runs initiated with a different *in silico* hit from the same Standard BinderFlow run. The grey dashed lines indicate the PAE interaction and pLDDT binder scores of the design obtained from the standard BinderFlow pipeline used as backbone to initialise Partial Diffusion and Sequence Diversity runs. The black, dotted lines indicate typical thresholds to consider a binder a hit, i.e., PAE_interaction < 10, pLDDT_binder > 80.

B) Correlation between PAE_interaction scores and Shape Complementarity scores for designs resulting from the Partial Diffusion run and the original candidate.

C) Correlation between PAE_interaction scores and Shape Complementarity scores for designs resulting from the Sequence Diversity run and the original candidate.

D) Correlation between PAE_interaction scores and BLOSUM62 scores, a measure of sequence similarity, calculated between binders resulting from the Sequence Diversity run and the original candidate.

E) Sequence alignment of binders obtained from a Sequence Diversity run. Top 20: the 20 higher-scoring binders ordered by PAE_interaction. Ref: the sequence of the binder used as template for initiating the Sequence Diversity run. Bottom 20: the 20 lower-scoring binders ordered by PAE_interaction.

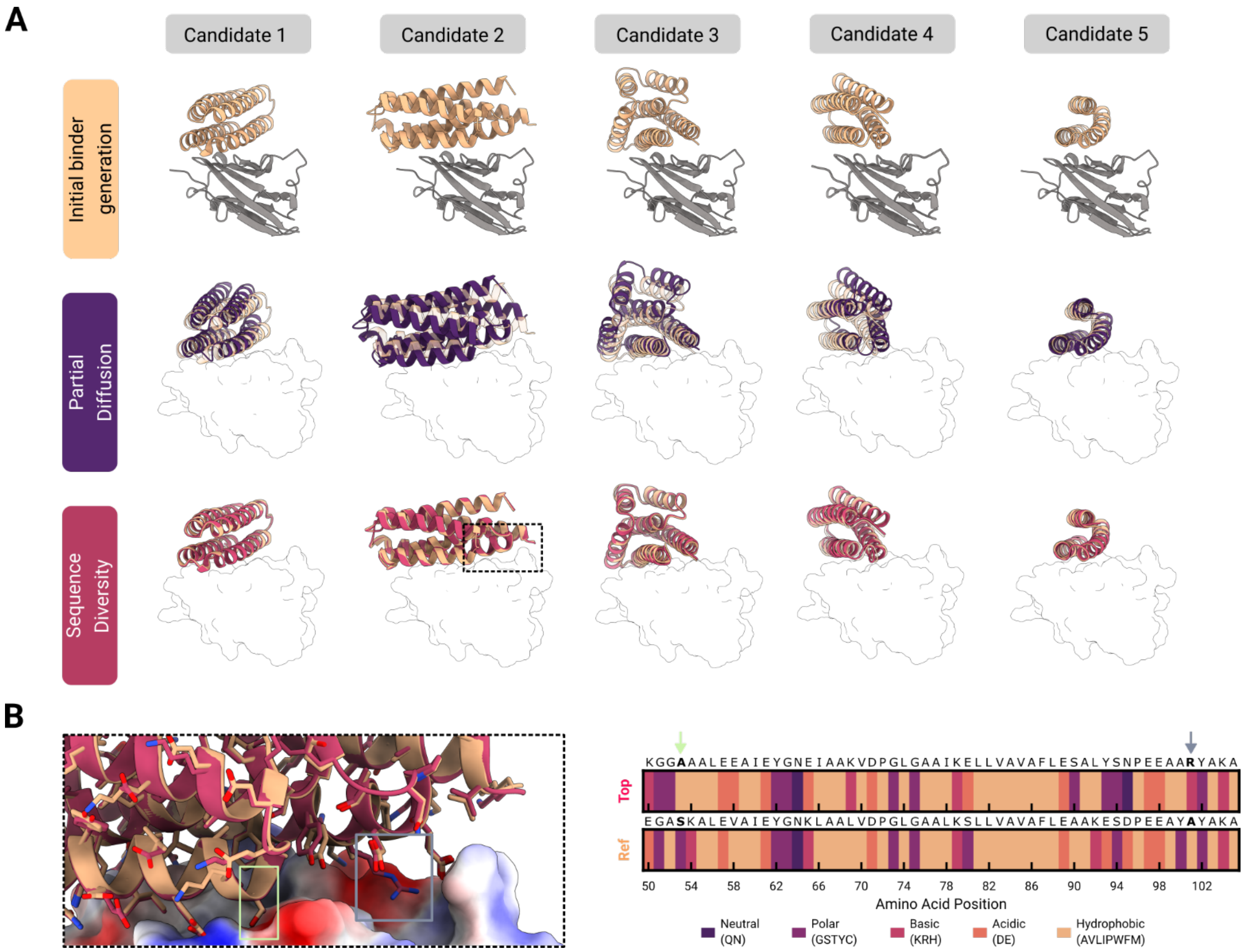


**Supplementary Fig. 2 – *In silico* refinement of candidates by Partial Diffusion and Sequence Diversity**

A) Structural alignments between candidates selected for *in silico* refinement (yellow) and example hits from Partial Diffusion (purple) and Sequence Diversity (red). The target, PDL1, is shown as a grey cartoon or as a silhouette. The first row displays the initial structure of the candidates, while the second and third rows show backbone structural variation for each candidate after the respective refinement strategy.

B) Residue identity changes introduced during Sequence Diversity. Left: *in silico* predictions of the input reference (yellow, Ref) and the top hit according to the PAE interaction (red, Top), with PDL1 shown as an electrostatic surface. Right: sequence alignment of the interacting helix (residues 50–105) between Ref and Top. Two residue changes are highlighted in lime (S > A) and grey (A > R).

**A**

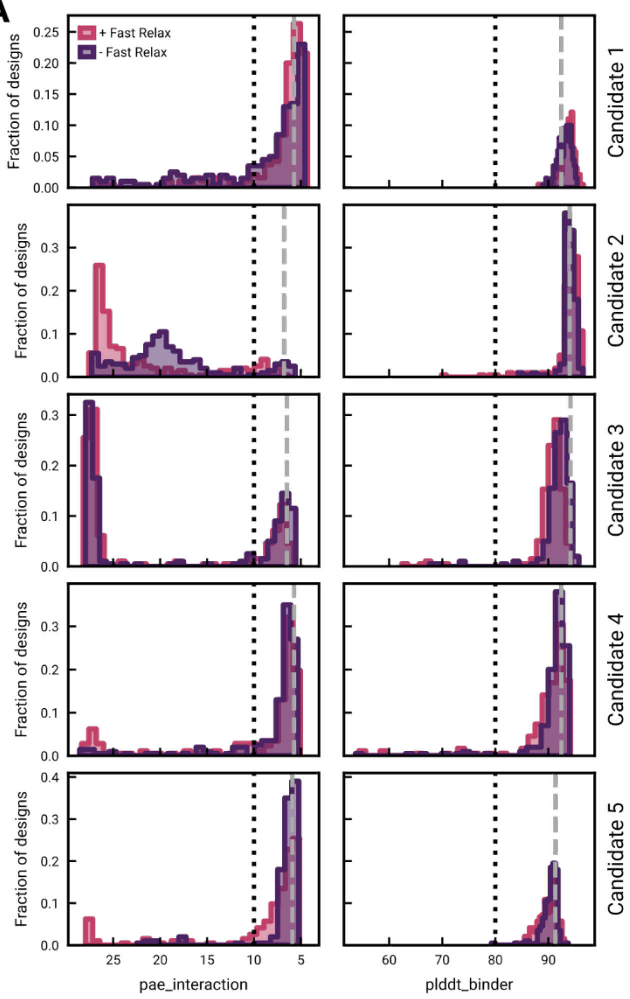


**B**

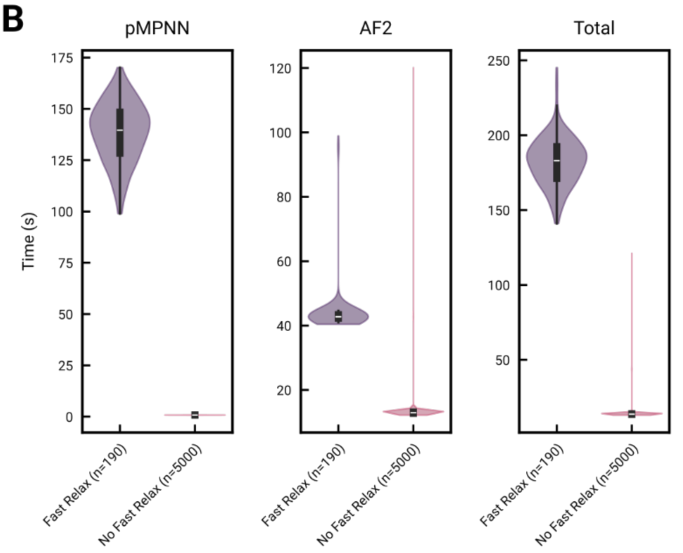


**C**

| Candidate run | Hit Rate + FR | Hit rate - FR |
|---|---|---|
| #1 | 73.03% (130/178) | 83.02% (914/1101) |
| #2 | 9.6% (19/198) | 13.76% (743/5401) |
| #3 | 30.96% (61/197) | 41.13% (2139/5201) |
| #4 | 85.86% (164/191) | 68.92% (3446/5000) |
| #5 | 80.93% (157/194) | 97.08% (5049/5201) |

**Supplementary Fig. 3 – FastRelax does not consistently improve Sequence Diversity *in silico* hit rates**

A) Comparison of PAE interaction and binder pLDDT distributions resulting from campaigns ran using Sequence Diversity with and without the FastRelax protocol. Each row represents runs initiated with a different *in silico* hit from the same Standard BinderFlow run. The black, dotted lines indicate typical thresholds to consider a binder a hit, i.e., PAE_interaction < 10, pLDDT_binder > 80.

B) Comparison of wall time per step in Sequence Diversity using, or not, the FastRelax. The number of designs corresponding to each step is indicated on the x-axis.

C) Comparison of *in silico* hit rates for five different Sequence Diversity runs with and without the FastRelax protocol. The raw count of hits for each candidate is indicated in parentheses.

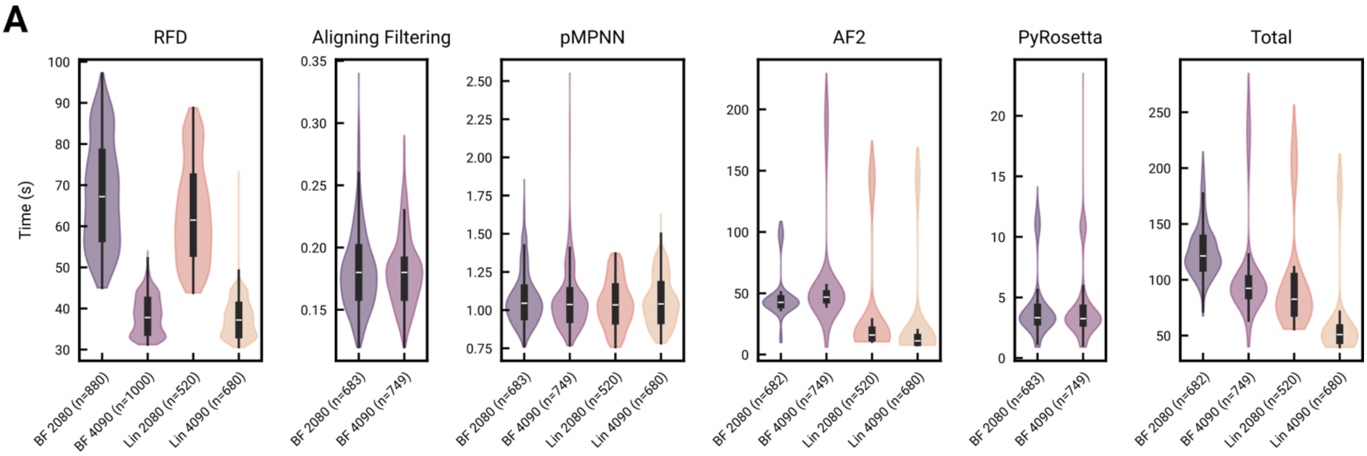


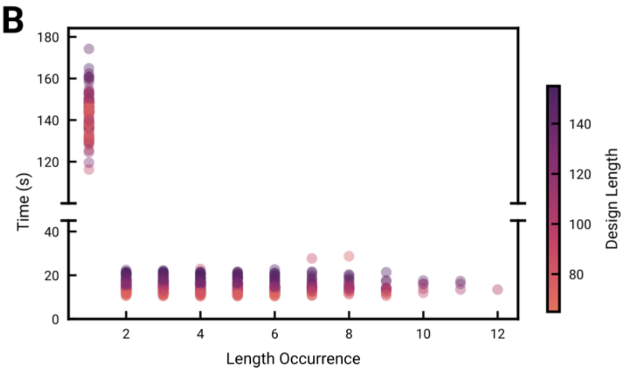


C

| | Linear (2080Ti) | BinderFlow (2080Ti) | Linear (4090Ti) | BinderFlow (4090Ti) |
|---|---|---|---|---|
| **RFD (s)** | 63.48 | 68.01 | 37.56 | 38.4 |
| **Filtering (s)** | - | 0.18 | - | 0.18 |
| **pMPNN (s)** | 1.02 | 1.08 | 1.08 | 1.08 |
| **AF2IG (s)** | 36.12 | 51.30 | 28.74 | 63.0 |
| **PyRosetta (s)** | - | 4.26 | - | 4.2 |
| **Total (s)** | **100.68** | **124.8** | **67.44** | **108.24** |

**Supplementary Fig. 4 – Benchmarking of BinderFlow**

A) Comparison of wall time per pipeline step between the linear approach (Lin) and BinderFlow (BF). The same campaigns were executed on RTX4090 and RTX2080 GPUs. The number of designs corresponding to each step is indicated on the x axis. “Aligning Filtering” and “PyRosetta” steps are only performed by BinderFlow.

B) Order of prediction impacts AF2IG scoring wall times. Length Occurrence stands for how many designs of that same length have been predicted in that same batch. Times extracted from the linear pipeline.

C) Table summarising the time spent per binder and design strategy in each step, comparing GPU models and binder generation pipelines.

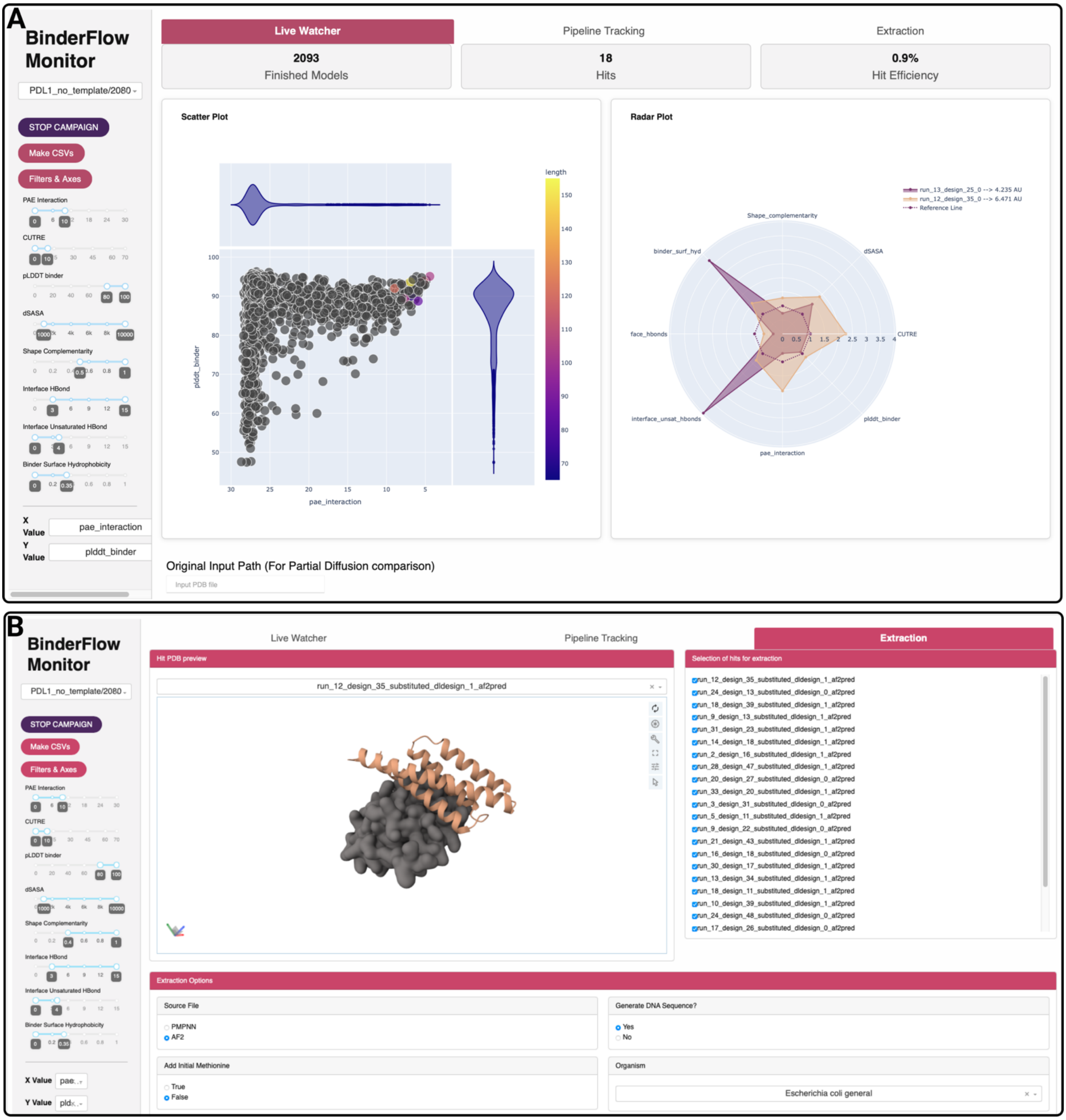


**Supplementary Fig. 5 – BFmonitor overview**

A) **Live Watcher** allows to monitor binder campaigns in real time and filter hits using more or less stringent thresholds for all the scores calculated by AF2IG and PyRosetta. These thresholds can be adjusted with the sliders on the left panel of the dashboard.

B) **Extraction** allows for visual inspection of the backbone of the selected hits and extraction of their structures as PDBs or their sequences as FASTAs. It also contains tools for codon optimized, reverse translation of the protein sequences, yielding ready-to-order DNA coding sequences.
